# Theranostic multifunctional lipid nanoparticles containing curcumin for integrated imaging and stabilizing vulnerable atherosclerotic plaques through an “eat-me” signal

**DOI:** 10.1101/2023.10.17.562822

**Authors:** Zhang Shi, Jun Huang, Chao Chen, Xuefeng Zhang, Zhiqiang Ma, Qi Liu

## Abstract

**BACKGROUND:** Curcumin has emerged as a promising candidate capable of polarization of macrophages, which promote the stability of atherosclerotic plaque. Nevertheless, a notable limitation lies in the non-specific nature of curcumin’s targeting. The present study endeavors to harness multifunctional lipid nanoparticles (MLNPs), which could both facilitate imaging and achieve targeted delivery of curcumin specifically to inflammatory macrophages, to effectively counteract vulnerable plaque and mitigate the risk of ischemic events.

**METHODS:** The term “MLNPs”, for targeted delivery of curcumin using multimodal imaging techniques including single photon emission computed tomography (SPECT) and magnetic resonance imaging (MRI), refers to a new type of nanoparticle designed to specifically target and modulate macrophages with phagocytic function. These nanoparticles are cholesteryl-9-carboxynonanoate-(^125^I-iron oxide nanoparticle/Cur)-lipid-coated nanoparticles [9-CCN-(^125^I-ION/Cur)-LNPs], which carry hybrid imaging agents. These agents are combinations of 125I-ION and lipids that contain phagocytic “eat-me” signals, which induces macrophages to swallow MLNPs.

**RESULTS:** The accumulation of the devised 9-CCN-(^125^I-ION/Cur)-LNPs on the unstable plaque of animal models in vivo was accurately reflected and lesions were highlighted by both SPECT and MRI. The intense radioactive signals on SPECT images facilitated the identification and quantification of the target lesion, while MRI based on ION particles facilitated the visualization of the focal localization and volumetry of atherosclerotic plaque. The targeted distribution of the unstable plaque in the rabbit aorta was further confirmed by ex vivo planar images of nuclide and Prussian blue staining for ION. Additionally, 9-CCN-(^125^I-ION/Cur)-LNPs were found to specifically and effectively bind to the pro-inflammatory M1 macrophages present in the unstable plaque, resulting in the accumulation of radionuclide and hypointensity on T2W images.

**CONCLUSIONS:** The 9-CCN-(^125^I-ION/Cur)-LNPs demonstrated remarkable capability in the delivery of both ^125^I-ION and curcumin to macrophages, ultimately resulting in M1-M2 macrophage polarization, which may offer valuable insights into addressing unstable atherosclerotic plaque.

## Introduction

The rupture of vulnerable atherosclerotic plaque can lead to acute ischemic events and infarction, imposing a significant risk of mortality and disability to individuals. Accurately predicting the progression and vulnerability of atherosclerotic plaque has However, achieving this precise prediction to conventional imaging modalities such as ultrasonography and angiography.

Recent studies have shed light on the pivotal role of macrophages in the initiation, development, and rupture of atherosclerotic plaque [1]. However, it is important to recognize that macrophages exhibit a heterogeneous nature, characterized by two distinct polarities: the pro-inflammatory “M1” macrophages and the anti-inflammatory “M2” macrophages [2,3]. Within this context, the “M1” macrophages demonstrate heightened phagocytic capability and play a significant role in the progressive advancement of atherosclerosis due to their active uptake of lipids and oxidized lipoproteins [4, 5]. Conversely, the “M2” macrophages efficiently phagocytose apoptotic cells, though they exhibit impaired phagocytic activity in the context of advancing atherosclerotic plaques [6,7]. As such, the modulation of macrophage behavior holds promising potential as an effective therapeutic strategy to address atherosclerosis.

Curcumin, a natural polyphenol extracted from Curcuma longa, has been identified as a potential modulator of the transition of pro-inflammatory M1 macrophages to the M2 phenotype. Its remarkable antioxidant, anti-inflammatory, anticoagulant, and lipid-lowering properties render it a promising candidate for targeting chronic inflammatory diseases. [8, 9]. Specifically, curcumin exerts its regulatory effects by inhibiting Toll-like receptor 4 (TLR4)-mediated mitogen-activated protein kinase (MAPK)/NF-κB pathways. This is achieved through downregulation of TLR4 and suppression of NF-κB and various MAPKs, including ERK (extracellular signal–regulated kinases), JNK (c-Jun N-terminal kinase), and p38 [7].

However, it is crucial to acknowledge that curcumin exhibits non-specific targeting to inflammatory macrophages. Therefore, an alternative strategy could be implemented that entails the development of a defined signaling pathway specifically designed to target these macrophages. To this end, the utilization of phosphatidylserine (PtdSer) and an oxidized cholesterol ester derivative known as cholesterol-9-carboxynonanoate (9-CCN) – both possessing “eat-me” signals – may efficiently enhance macrophage phagocytosis [10]. By capitalizing on this targeted strategy, the risk of plaque rupture could potentially be mitigated through the selective delivery of curcumin to inflammatory macrophages, thereby facilitating the transformation of macrophages from the M1 to the M2 phenotype.

In recent times, magnetic iron oxide nanoparticles (IONs) have received significant attention for the detection of atherosclerotic plaque, primarily due to their unique hypointensity appearance on T2-weighted (T2W) magnetic resonance imaging (MRI) [11–13]. To minimize the potential adverse impact of IONs on human subjects, we adopt a clinical strategy that involves pooling IONs within ligand-conjugated nanomedicines, including nanoliposomes and biodegradable polymer nanoparticles[13–16]. Despite these advancements, the sensitivity and signal-to-noise ratio (SNR) of T2W images in conventional magnetic fields limiting factors, hindering the ability of IONs to provide accurate imaging of the cardiovascular system [17]. To circumvent this limitation, a multimodal imaging approach has been developed that involves the integration of two or more different imaging modalities, such as MRI with nuclear medical imaging, computed tomography (CT) imaging, or fluorescence imaging. This approach has shown potential in enhancing imaging specificity and resolution [11, 13].

As a vivid illustration, traditional MRI typically requires the presence of >10^7^ Gd atoms [18], or a high relaxivity of >10^6^ for IONs [19] to detect a single cell. In stark contrast, SPECT-PET imaging achieves this feat with only approximately 0.01–0.2 radio atoms per cell [18]. In our previous study, we successfully combined MRI and SPECT by labeling 99mTc onto the surface of IONs. This innovative approach provided highly complementary information for in vivo characterization of atherosclerotic plaque [20], holding great promise in advancing our understanding of atherosclerosis and potentially enabling more precise and comprehensive plaque assessment.

The overarching goal of the current investigation is to engineer a theranostic lipid-coated nanoparticle that can jointly deliver a combination of imaging agents, specifically ^125^I-IONs and curcumin, with specific affinity towards inflammatory macrophages within the plaque, facilitated by the incorporation of “eat-me” signals (PtdSer and 9-CCN) within the lipid membrane. Once internalized by the targeted macrophages, curcumin is released into the cytoplasm, while the hybrid imaging agents are engulfed. This process can subsequently be detected through the observation of low signal intensity on T2W images, and the signal emitted by radioactive elements using SPECT. The development of such a multifunctional nanoparticle holds great promise for enhancing our ability to therapeutically and diagnostically address atherosclerotic plaque and its associated complications.

## 1. Materials and methods

### 1.1. Materials

1,2-distearoyl-sn-glycero-3-phosphoethanolamine-N-[methoxy(polyethylene glycol)-2000] (DSPE-PEG), L-α-phosphatidylcholine and DOPS were purchased from Avanti Polar Lipids (Alabaster, AL). 9-CCN was synthesized in supplementary information as our previous work.[10] Na^125^I was purchased from Shanghai GMS Pharmaceutical Co., Ltd. (Shanghai, China). Streptavidin-functionalized IONs (10 nm), curcumin, and iodogen (1,3,4,6-tetrachloro-3α,6α-diphenylglucoluril) were purchased from Millipore-Sigma (Billerica, MA, USA). All other reagents were of analytical grade and purchased from Sinopharm (Shanghai, China).

The macrophage cell line, RAW264.7, was obtained from Shanghai CAS (Chinese Academy of Sciences, Shanghai, China). The cells were cultured in Dulbecco’s modified Eagle’s medium (Gibco, Grand Island, NY, USA) supplemented with 10% (vol/vol) FBS (Gibco, Grand Island, NY, USA), 100 U/mL penicillin, and 100 μg/ml streptomycin at 37 ℃ in a humidified atmosphere of 5% CO_2_/95% air. Human umbilical vein endothelial cells (HUVECs) were purchased from Cascade Biologics (Portland, OR, USA), and cultured in Medium 200 supplemented with Low Serum Growth Supplement (Gibco, Grand Island, NY, USA) in a 37 ℃, 5% CO_2_/95% air, humidified cell culture incubator.

### 1.2. Fabrication and characterization of nanoparticles

The multifunctional lipid nanoparticles for imaging and targeted delivery of curcumin was fabricated and characterized as describing in supplementary information, partially according to our previous research[13].

### 1.3. Modulating and imaging macrophages from M1 to M2 in vitro

#### 1.3.1. Uptake of nanoparticles in macrophages and HUVECs

RAW264.7 cells and HUVECs (2.5 × 10^5^ cells per well) were seeded overnight on coverslips in a 12-well tissue culture plate and treated with and (^125^I-ION/Cur)-LNPs for 2 h (curcumin concentrations: 50 μg/mL). After washing with PBST (PBS containing 0.1% Tween-20), the cells were fixed with 4% paraformaldehyde. Cells were then incubated with 40,6-Diamidino-2-phenylindole dihydrochloride (Sigma, Fluka Chemie, Buchs, Israel) for nuclear staining, and mounted under glass coverslips. Immunofluorescence of the cells was visualized under a Confocal LSM510 microscope (Carl Zeiss Microimaging, Inc., Thornwood, NY).

Quantitative analysis of curcumin uptake in RAW264.7 and HUVECs was conducted using the HPLC assay. After overnight inoculation of RAW264.7 cells (5 × 10^5^ cells per well) in 12-well plates, the cell medium was replaced with fresh medium containing MLNPs and (^125^I-ION/Cur)-LNPs (curcumin concentrations: 50 μg/mL). After incubation for another 24 h, the cells were collected after enzyme-digesting, sonicated, and the supernatant containing curcumin after centrifugation was measured by HPLC as described above.

The cell lines were imaged on SPECT/CT and MRI to evaluate the binding reactivity of nanoparticles to RAW264.7 and HUVECs. Briefly, 1 × 10^6^ of RAW264.7 and HUVEC cells were vortexing incubated with 2 kBq of nanoparticles in growth medium for 4-h at 37 ℃. After washing in PBS, the cells were homogenously suspended in 0.8 ml of 2.5% agar gel in a 24-well plate. Planar gamma imaging was performed on a clinical SPECT/CT scanner (Precedence, Philips Healthcare) equipped with a low-energy, high-resolution parallel-hole collimator. The radioactivity nanoparticles in the above cell lines were also quantitatively counted using a GC-1200 γ counter (ZONKIA, Hefei, China). Then, MR imaging was performed on a clinical MRI scanner (Siemens 3.0T) equipped with a microsurface coil using a spin echo sequence. T2WI parameters were as follows: matrix = 256×256, FOV = 12 × 12 cm, section thickness = 1 mm, TE = 45 ms, TR = 3000 ms, number of excitations (NEX) = 1. Furthermore, the same cell samples were further lysed and digested with 2% HNO_3_ for intracellular iron content quantification using an ICP-AES instrument (Perkin Elmer Optima 8000).

#### 1.3.2. Reverse Transcription and Real-time PCR

RAW 264.7 macrophages (5 × 10^5^ cells/well) were seeded in a 6-well transwell plate and incubated overnight. The cell medium was then replaced with fresh medium containing nanoparticles or curcumin at a concentration of 5 μg/mL. After 48 h, the cells were analyzed for the mRNA expression of Arginase 1 (Arg-1), macrophage galactose-type C lectin 1 (Mgl1), and inducible nitric oxide synthase (iNOS) as described in supplementary information.

#### 1.3.3. Enzyme-Linked Immunosorbent Assay (ELISA)

RAW 264.7 macrophages (5 × 10^5^ cells/well) were seeded in a 6-well transwell plate and incubated overnight. The cell medium was then replaced with fresh medium containing nanoparticles or curcumin at a curcumin concentration of 5 μg/mL. After 48 h, cell culture supernatants were harvested and analyzed. The secretion of TNF-α, IL10, and IL12 was assayed using an ELISA kit (Boster Biological Technology, Wuhan, China). All experiments were conducted in triplicate.

#### 1.3.4. Flow cytometry for CD206 and CD16/32 expression measurement

RAW 264.7 macrophages were stimulated by IFN-γ (20 ng/mL) and lipopolysaccharide (LPS, 100 ng/mL) for 12 h, detached from the plates using trypsin, and washed with PBS. Cell suspension (100 μL) was aliquoted into 1-mL Eppendorf (EP) tubes. Then, double antibody staining was performed as follows: anti-mouse CD206 PE (Biolegend, CA, USA) and anti-mouse F4/80 FITC (Biolegend, CA, USA), or anti-mouse CD16/32 PE (Biolegend, CA, USA) and anti-mouse F4/80 FITC (Biolegend, CA, USA) were added to the cells and incubated for 0.5h at 4 ℃. The cells were then washed twice with PBS and resuspended in 300 μL 1 × PBS solution before analysis on Becton-Dickinson FACS Calibur (Becton-Dickinson, Bedford, MA).

### 1.4. Modulating and imaging macrophages from M1 to M2 in vivo

#### 1.4.1. Macrophage isolation from the atherosclerotic plaque in the thoracoabdominal aorta of rabbits

Macrophages were isolated from the plaque in the thoracoabdominal aorta of the rabbits as described in supplementary information, similar to previous protocol.[4] The isolated macrophages were analyzed by flow cytometry to examine the M1/M2 proportion with anti-mouse CD206 FITC and anti-mouse CD16/32 PE as described above. Furthermore, the cells were also analyzed for the mRNA expression of Arg-1, Mgl1, and iNOS as described above. To analyze the cytokines of TAM, the isolated TAM was fixed, permeated, and stained with TNFα-PE, IL10-PE, and IL12-PE (BD BioSciences, USA) for 30 min at 4 ℃ away from light. After washing with PBS twice, the samples were detected using a flow cytometer (BD FACS VERSE; BD BioSciences USA) for quantitative analysis.

#### 1.4.2. Rabbit model of the atherosclerosis and ^125^I-ION Hybrid images

The New Zealand white rabbit model of atherosclerosis (male, 2.5∼3 kg) was established by abdominal aorta balloon injury plus a high-cholesterol diet (80% standard feed + 1% cholesterol + 5% lard + 10% egg yolk powder) for 10 weeks. After injection of ^125^I-ION via the auricular veins (one doses, 12.3 MBq/kg, ∼1.67 mg iron/kg), the rabbits were anesthetized via 2% isoflurane inhalation and imaged consecutively with clinical SPECT/CT scanner (Siemens symbia T16 SPECT/CT) and a 3.0 T clinical MRI scanner (Signa HDxt, GE Healthcare, USA) at 6 and 36 h post-injection. CT was performed with the following scan parameters: frame resolution, 512 × 512; tube voltage, 140 kV; current, 0.15 mA; exposure time, 500 ms/frame. Each scan took approximately 7 min. Real-time 3D reconstruction of the collected images was performed using Nucline Software (v1.02, Mediso, Budapest, Hungary). SPECT was performed after CT scanning using the following parameters: collimator, low-energy high-resolution (LEHR), energy peak, 35 keV; window width, 20%; matrix, 128 × 128; scan time, and 45 s/projection with 32 projections in all. The whole-body scan for each rabbit took 24 min on average, and three-dimensional ordered-subset expectation maximization images were reconstructed. At 36 h after injection, the thoracoabdominal aorta of the rabbits was examined by MRI fast spin-echo (FSE) T_2_W imaging using a 3.0 T MR scanner (Signa HDxt, GE Healthcare, USA) and an 8-channel phased abdominal coil. The MR parameters were as follows: matrix = 320 × 256, FOV = 13 × 13 cm, section thickness = 2 mm, TE = 51 ms, TR = 2884 ms, NEX = 3. The atherosclerosis in the thoracoabdominal aorta of the rabbits was further confirmed by the paraffin sections and H&E staining. The macrophages in the atherosclerosis in the thoracoabdominal aorta of the rabbits were revealed using mouse anti-rabbit CD68 antibody (Abcam, Cambridge, UK) as the primary antibody and goat anti-mouse IgG antibody (horseradish peroxidase (HRP) labeled) as the secondary antibody.

#### 1.4.3. In vivo MRI and SPECT imaging of atherosclerotic plaques

New Zealand white rabbits (male, ∼3 kg) with atherosclerosis were randomly administered with 9-CCN[^125^I-ION/Cur]-NPs (three doses, 12.3 MBq/kg, ∼1.67 mg iron/kg, ∼17 mg curcumin/kg, 6 rabbits per group). To verify the in vivo therapeutic efficacy, six rabbits were administered [^125^I-ION/Cur]-NPs (12.3 MBq/kg, ∼1.67 mg iron/kg). All nanoparticles were injected via the auricular veins once every 10 d for 30 d, according to the following investigations. The rabbits were imaged, and the atherosclerotic plaque was monitored by SPECT and MRI scanning at 6 and 36 h post-injection, respectively. The parameters were the same as those for ^125^I-ION SPECT and MRI images.

The SPECT transverse images are slices from the 3D reconstruction with a section thickness of 3 mm. The SPECT image segmentation and quantitation analysis were performed using the InVivoScope 1.40 software. Briefly, the volume of interest (VOIs) was obtained in the form of a cylinder by first circling the ROIs from the transverse profile, and then selecting the length of ROIs from the maximum intensity projection. Following the above operation, the values of radioactivity and volume of the VOIs were obtained from the software. The radiotracer uptake (Bq/voxel) was calculated by dividing radioactivity by volume from VOIs. For MRI quantitative analysis, we calculated the SNR in the same three regions of interest (ROI, 20 × 20 pixels) placed in the aortic wall of the lower intrathoracic aorta on three different coronal reformatted images[5].

#### 1.4.4. Ex vivo evaluation of atherosclerotic plaques

After the third MRI scan, rabbits were anesthetized and then sacrificed. A total of 100 mL of 0.9% NaCl and 20 ml of 4% paraformaldehyde were orderly and slowly injected into the heart to clean up the blood and fix the vessels. The aorta was carefully and completely dissected from the body for analysis. For *ex vivo* planar imaging of *in vitro* nuclide distribution, the aorta was placed in the central position of the SPECT camera (Precedence, Philips Healthcare) for planar imaging with the following parameters: collimator: LEHR, energy peak, 35 keV; window width, 20%; matrix, 128 × 128; scan time, 60 min. The iron distribution in the plaque was revealed by Prussian blue staining. H&E staining further confirmed AS in the thoracoabdominal aorta of the rabbits. The macrophages in the plaque in the thoracoabdominal aorta of the rabbits were revealed using mouse anti-rabbit CD16/32 and CD206 antibody (Abcam, Cambridge, UK) as the primary antibody and goat anti-mouse IgG antibody (HRP labeled) as the secondary antibody. The macrophages of atherosclerotic plaque in the thoracoabdominal aorta of the rabbits were also revealed using FITC-labeled mouse anti-rabbit CD16/32 and PE-labeled mouse anti-rabbit CD206 antibody (Abcam, Cambridge, UK). The organs of the rabbits were excised, and H&E staining was performed on the organs to examine the toxicity of the nanoparticles. The rabbits were weighed once per day for 4–5 days after drug treatment.

#### 1.4.5. Animal study

All rabbits were purchased from the Shanghai Experimental Animal Center of Chinese Academy of Sciences (Shanghai, China). All procedures were per formed in accordance with the guidelines of the Committee on Animals of the Naval Medical University (Shanghai, China).

### 1.5. Statistical analysis

Data were analyzed using the statistical package SPSS 24.0 (IBM, USA). For values that were normally distributed, a direct comparison between two groups was conducted by Student’s non-paired t test, and one-way analysis of variance (ANOVA) with the Dunnett’s or Newman Keuls post-tests were used to compare the means of three or more groups. A *p* value of < 0.05 was considered statistically significant.

## 2. Results

### 2.1. Construction and characterization of nanoparticles

The nanoparticles were synthesized using a one-step self-assembly method subsequent to the hydration of a lipid film with 4% aqueous streptavidin-functionalized ION (10-nm diameter), followed by rapid bath sonication. Each lipid film utilized a composition comprising 2–5 mol% of PtdSer and 9-CCN, 5 mol% of phosphatidylethanolamine-PEG2000, as well as varying concentrations of contrast agents, drugs, and dyes, as detailed in the following section and Supporting Information. Notably, we observed that the lipid film can accommodate up to 7 mol% of hydrophobic entities in any combination, thereby enabling the precise synthesis of particles with desired functionality (Figure 1).

**Figure 1.**
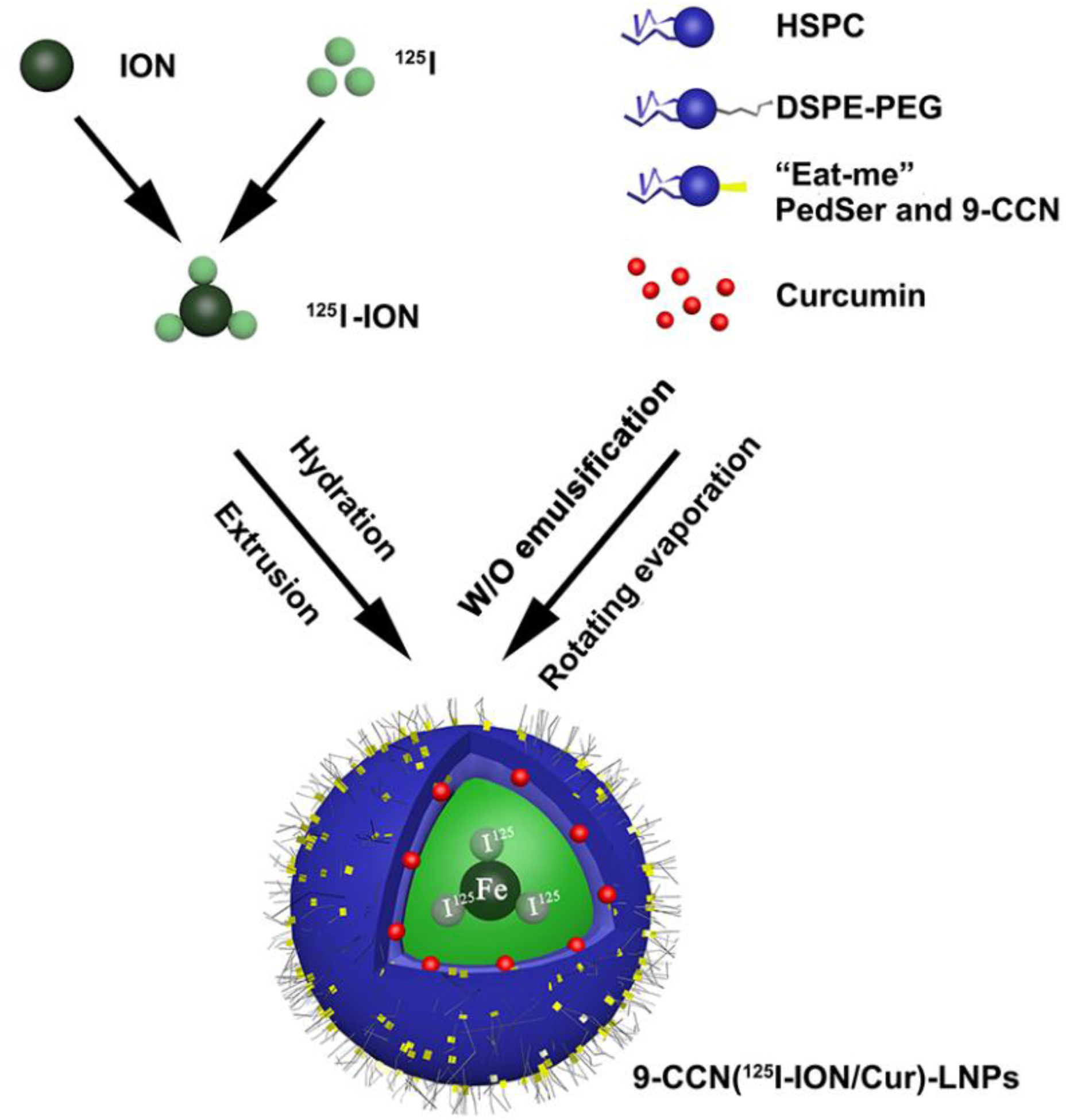
Synthesis route of lipid-coated ^125^I-IONs targeting macrophages (MLNPs).

The lipid-coated ^125^I-IONs with curcumin, denoted as MLNPs, were initially synthesized using the iodogen method. Subsequently, the MLNPs nanoparticles were prepared through the coating of ^125^I-IONs with a lipid film loaded with the hydrophobic drug curcumin. As a result, the synthesized MLNPs nanoparticles exhibit multifunctionality, encompassing macrophage targeting, MRI capabilities through the incorporated IONs, SPECT imaging facilitated by the radionuclide, and therapeutic effects attributed to the presence of curcumin.

The nanoparticle characteristics have been comprehensively summarized in Table 1. The size of the MLNPs, 9-CCN(^125^I-ION)-LNPs, and (^125^I-ION/Cur)-LNPs measured approximately 120 nm, exhibiting a negative zeta potential and a narrow polydispersity index (PDI) below 0.2. Encapsulation efficacy of curcumin (CEE) in MLNPs surpassed 86.4%, while the encapsulation efficacy of iron (FEE) was determined to be 32.5%. The transmission electron microscopy (TEM) images displayed moderately spherical and uniform shapes for both MLNPs and 9-CCN(^125^I-ION)-LNPS (Figure 2A). The relaxation rate, R2 = 1/T2, exhibited a linear relationship with the iron concentration (Figure 2B). The slopes obtained from the plot of R2 against iron concentration demonstrated comparable T2 relativities for MLNPs, ICC, and ^125^I-IONs (528.67, 540.66, and 554.25 s−1mM−1, respectively), affirming that the lipid coating of IONs does not attenuate their inherent superparamagnetic properties. Furthermore, the nanoparticle size remained stable throughout the 4-day study period, indicating favorable nanoparticle stability (Figure 2C). The in vitro release of curcumin from MLNPs was assessed in PBS and PBS + 10% fetal bovine serum (FBS) (Figure 2D). The release profile exhibited faster release kinetics in PBS + 10% FBS compared to PBS alone. Notably, a rapid release of curcumin from both nanoparticles was observed within the initial 30 hours, with over 40% released during this period. The cumulative release reached approximately 70% in PBS and approximately 85% in PBS + 10% FBS over the entire 240-hour duration.

**Figure 2.**
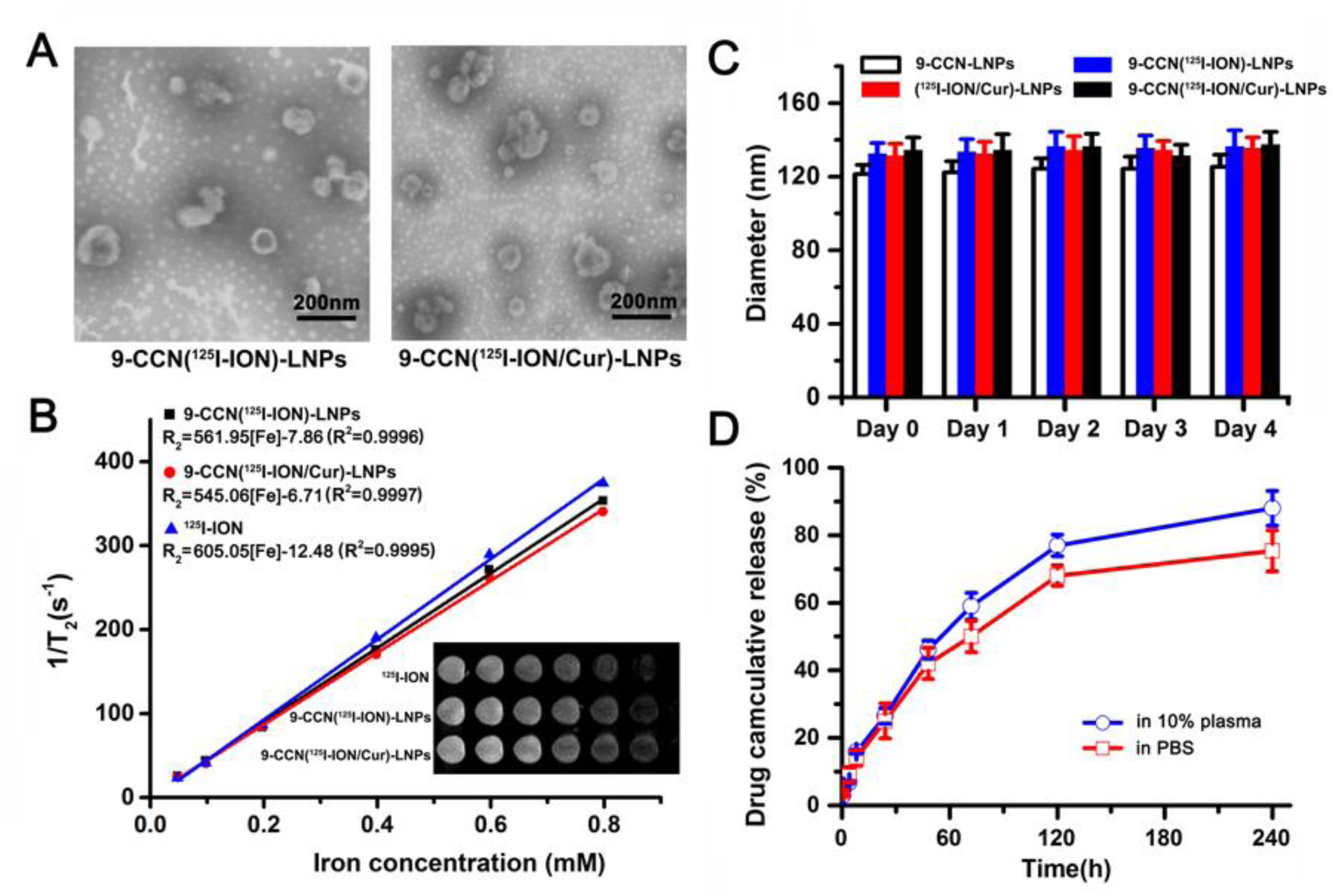
Characterization of nanoparticles. (A) The morphology under TEM. Bars represent 200 nm. (B) The NMR relaxometry of magnetic nanoparticles. The T2 relativity was calculated by a linear fit of the inverse relaxation times as a function of the iron concentration. (C) The stability of the nanoparticles in PBS with 10% FBS in 120 h. (D) The drug release of the nanoparticles in a period of 240 h. Data are expressed as mean ±SD (n = 3).

### 2.2. In vitro cellular uptake of nanoparticles

The internalization of nanoparticles was assessed using confocal microscopy. RAW264.7 and HUVECs were exposed to the nanoparticles for 2 hours at 37 ℃. The obtained results are depicted in Figure 3A, where it can be observed that RAW264.7 cells treated with MLNPs exhibited evident internalization of nanoparticles, manifested by the presence of green fluorescence indicative of curcumin. Conversely, RAW264.7 cells treated with ICC demonstrated no significant uptake or internalization, as denoted by the absence of discernible green fluorescence. When the nanoparticles were incubated with HUVECs, only minimal intracellular fluorescence was observed, suggesting a lack of cellular uptake for both MLNPs and ICC. These findings strongly suggested that MLNPs possesses specific targeting capabilities towards RAW264.7 cells, confirming the pivotal role of the “eat-me” signals, namely 9-CCN and 1,2-dioleoyl-sn-glycero-3-phospho-L-serine (DOPS), in facilitating the uptake of MLNPs in RAW264.7 cells. The intracellular uptake of curcumin in both RAW264.7 cells and HUVECs was quantitatively assessed to measure the quantity of internalized curcumin (Figure 3B). In accordance with the fluorescence-based assay, the curcumin concentration in MLNPs was significantly higher than that in ICC for RAW264.7 cells (P<0.05), but not in HUVECs, indicating that the eat-me signals of 9-CCN and DOPS effectively mediated the cellular uptake of curcumin in RAW264.7 cells (Figure 3B).

**Figure 3.**
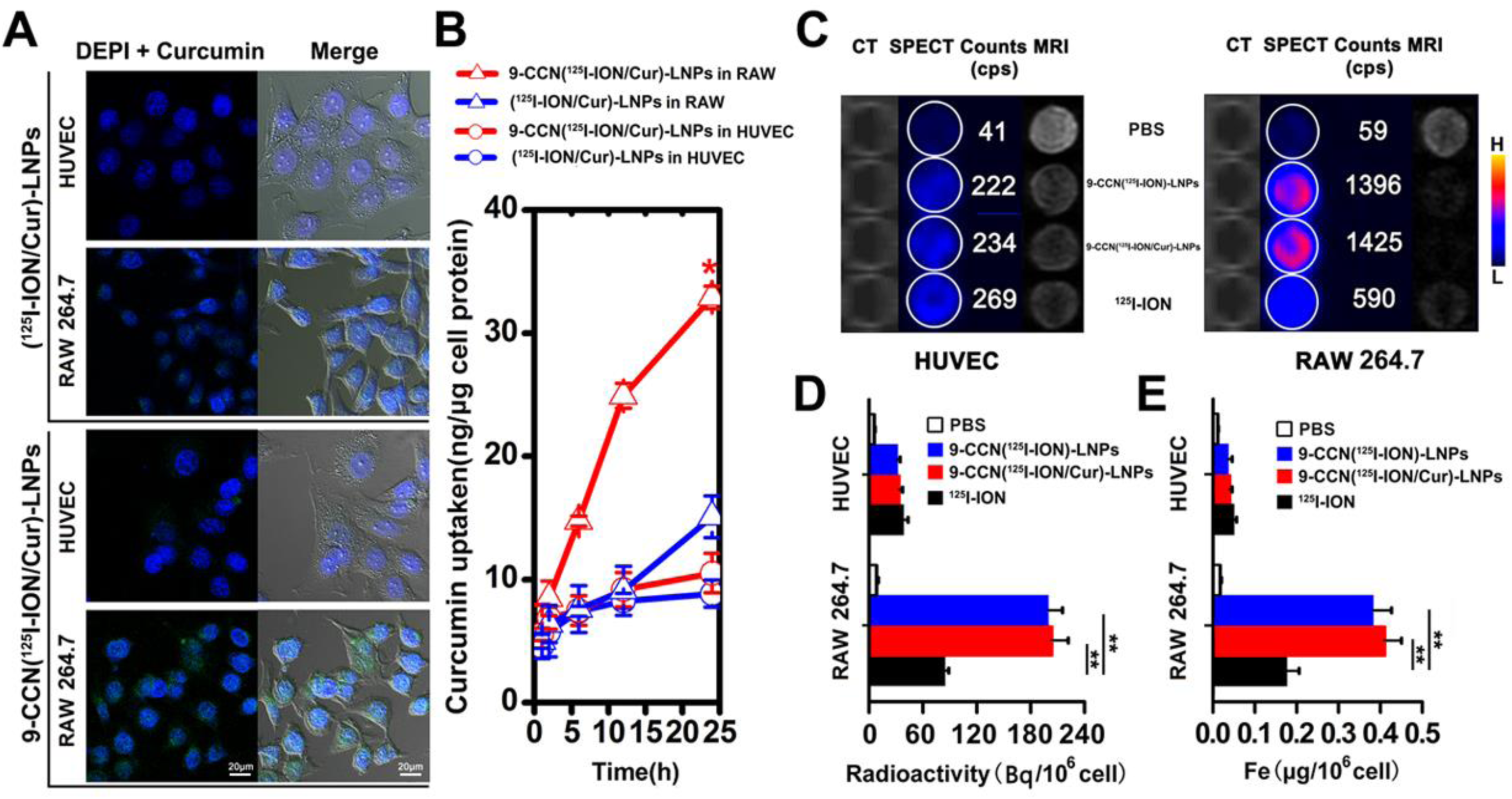
*In vitro* characterization of various nanoparticles in different cell lines. (A) Confocal microscopy evaluation of nanoparticles (green) uptake in macrophages (RAW 264.7 cells), HUVEC. The right panel represents the overlay of fluorescent and bright field (DIC) micrographs. (B) Time-dependent uptake of curcumin by different cell lines. (C) Planar gamma imaging performed on a clinical SPECT/CT scanner equipped with a low-energy, high-resolution parallel-hole collimator. (D) and (E) The radioactivity and intracellular iron content quantification of nanoparticles in the above cell lines. The three or more groups are compared by one-way ANOVA with Dunnett’s post-test. *, p < 0.05; **, p < 0.01; ***, p < 0.001; n.s. represents not significant (p > = 0.05). Data are expressed as mean ±SD (n = 3).

Furthermore, RAW264.7 and HUVECs incubated with 2 kBq 9-CCN[^125^I-ION/Cur]-NPs for 1 hour were readily detected using both SPECT and MRI. As shown in Figure 3C, the SPECT signal of 9-CCN[^125^I-ION]-NPs, with or without curcumin, was amplified by more than six-fold in RAW264.7 cells compared to that in HUVECs, and by more than two-fold compared with the non-targeted ^125^I-ION group in RAW264.7 cells. Similarly, MRI images exhibited the same trend in T2WI signals for both cell lines, indicating the specific binding of ION to M1 macrophages.

After PBS washing, the radioactivity of the incubated cells was measured using a gamma counter, and the internalized iron content within the cells was determined using ICP-AES. Following a 1-hour incubation of 9-CCN[^125^I-ION]-NPs, RAW264.7 cells displayed a radioactivity of 203.6 Bq, with 0.366 μg of iron internalized per 106 cells (Figure 3D). In HUVECs, the radioactivity was significantly decreased to 33.4 Bq, and iron uptake was also decreased (∼0.038 μg of iron per 10^6^ HUVECs). The quantitative assay was in accordance with the SPECT and MRI results.

### 2.3. MLNPs induces M2 macrophage polarization in vitro

Figure 4A presents the in vitro demonstration of M2 macrophage polarization. Flow cytometry analysis revealed a notable increase in the proportion of M2 macrophages (CD206+/F4/80+) in both MLNPs and curcumin-treated RAW264.7 cells. To delve into the intricate molecular alterations associated with macrophage polarization in vitro, reverse transcription-polymerase chain reaction (RT-PCR) was employed to assess the expression differences of M1-versus M2-type mRNAs. The mRNA levels of Arg1, Mgl1, and iNOS were quantified. As depicted in Figure 4B, macrophages exposed to MLNPs and curcumin displayed significantly heightened expression of M2-associated Arg1 and Mgl1, along with reduced expression of iNOS, in comparison to 9-CCN(^125^I-ION)-LNPS and PBS-treated macrophages. Additionally, enzyme-linked immunosorbent assay (ELISA) was conducted to investigate the secretion of IL-10, IL-12, and TNFα. The results illustrated in Fig. 4C demonstrated a substantial increase in the production of IL-10 and a decrease in the production of IL-12 and TNFα. These findings collectively confirm that MLNPs and curcumin effectively induced macrophage polarization towards an M2 phenotype.

**Figure 4.**
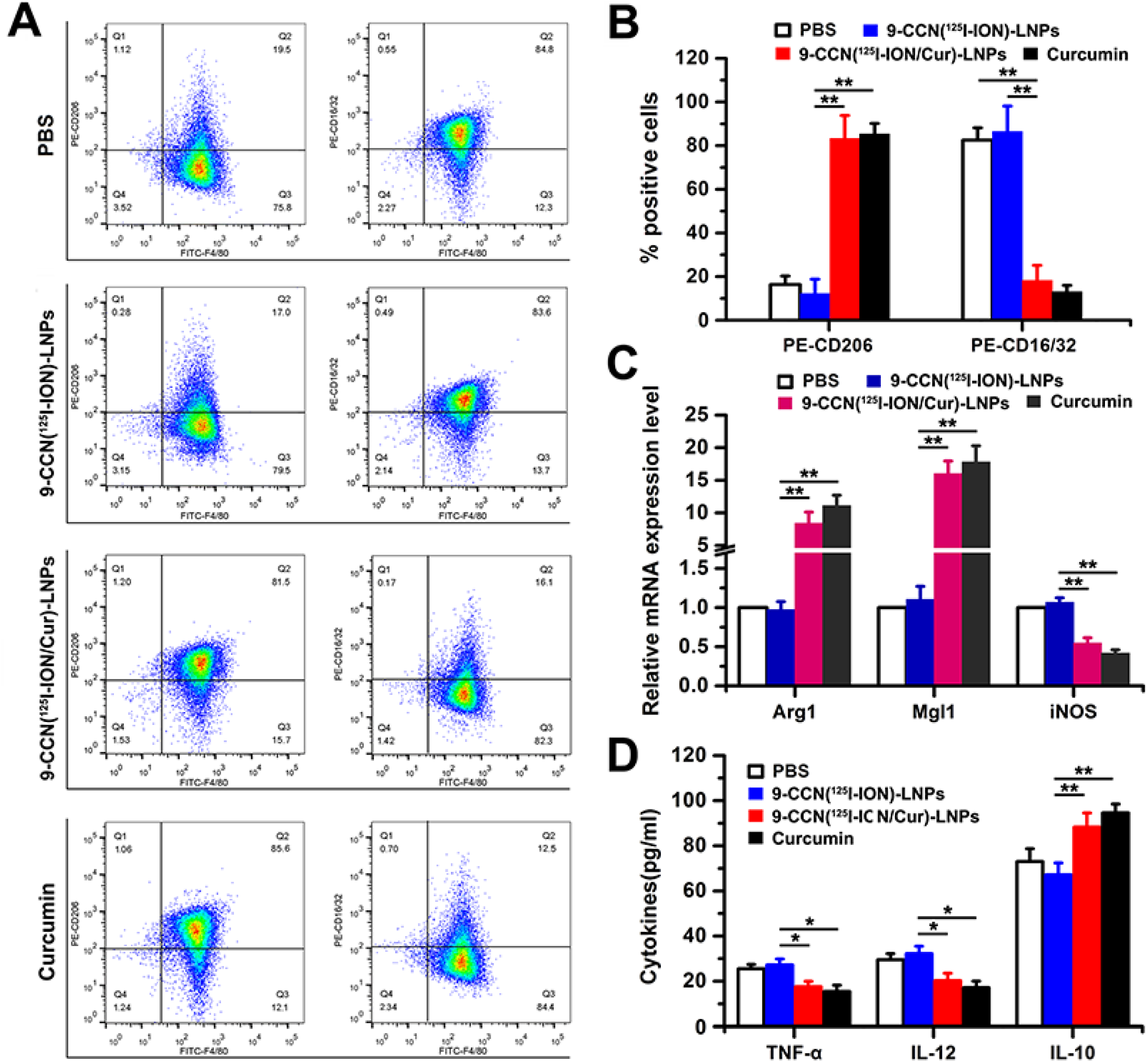
The M1-M2 polarization of RAW264.7 macrophages. The analysis of CD206, F4/80 and CD16/32 expression of macrophage by flow cytometry. (A) The proportion of positively stained cells. A representative experiment was shown from three independent experiments. (B) The quantitative analysis of CD206+ and CD16/32+ macrophages in (A). (C) The mRNA levels of M1 and M2 macrophage markers (Arg1, Mgl1 and iNOS) and (D) the cytokines concentrations (TNF-α, IL-12 and IL-10) in RAW264.7 macrophages. The three or more groups are compared by one-way ANOVA with Dunnett’s post-test. *, p < 0.05; **, p < 0.01; ***, p < 0.001; n.s. represents not significant (p > = 0.05). Data are expressed as mean ±SD (n = 3).

### 2.4. MLNPs visualizes atherosclerosis model by SPECT and MRI

Initially, in the healthy rabbits injected with ^125^I-ION, SPECT images indicated an absence of detectable ^125^I signals (Figure 5A), and the MR images of the ascending and abdominal aorta exhibited no noticeable T2WI signals (Figure 5B). In stark contrast, as depicted in Figure 5C, the ^125^I signal originating from the atherosclerotic plaque in the ascending aorta became evident at 6 hours post-injection, concomitant with an increase in signals from the heart, lung, and bladder during the initial 6-hour period, as observed by SPECT (Figure 5C). Accordingly, sagittal views of 3D time-of-flight magnetic resonance angiography (3D-TOF-MRA) demonstrated concurrence with mild stenosis at the same location, with transverse views of the corresponding slice exhibiting a notable hypointensity in T2WI along the thoracic and abdominal aorta due to the accumulation of IONs within the atherosclerotic plaque at 36 hours post-injection (Figure 5D&E). Subsequently, immunohistochemistry tests confirmed the presence of macrophages in the highlighted lesions, evident by the abundantly stained cell nuclei of the thoracic and abdominal aorta in the CD-68 stained plaques (Figure 5F&G). These results conclusively validate the successful establishment of the atherosclerotic rabbit model and demonstrate the accurate localization and detection of the plaque using SPECT and MRI, based on the aforementioned parameter settings.

**Figure 5.**
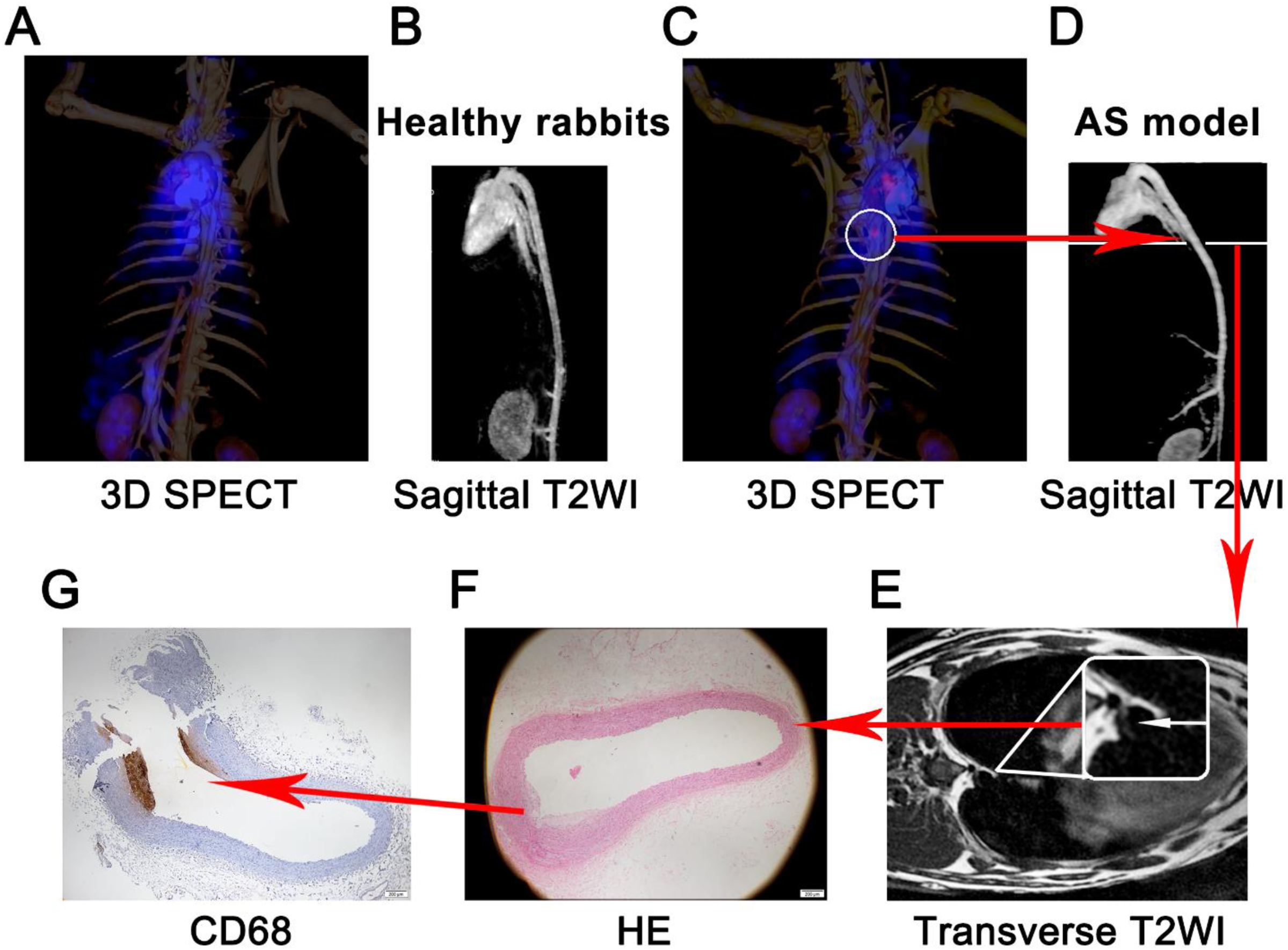
The 3D SPECT and T2-weighted imaging (T2WI) of healthy rabbits and AS model of rabbits. (A) The 3D SPECT of healthy rabbits. (B) The sagittal T2WI of healthy rabbits. (C) The 3D SPECT of AS model of rabbits. (D) The sagittal T2WI of AS model of rabbits. (E) The transverse T2WI of AS model of rabbits. (F) The AS in the thoracoabdominal aorta of the rabbits further confirmed by the paraffin sections and H&E. (G) The macrophages in the AS in the thoracoabdominal aorta of the rabbits.

### 2.5. MLNPs detects and counters atherosclerosis in vivo

The in vivo evaluation of nanoparticle MR and SPECT imaging was conducted in rabbits with atherosclerosis, as depicted in Figure 6. Advanced lesions were confirmed by 9-CCN[^125^I-ION]-NPs, both with and without curcumin, wherein radionuclides were predominantly localized in the ascending aorta at 6 hours post-injection. Simultaneously, significant accumulation was observed in the T2W image of the same slice at 36 hours post-injection. The integration of these MR images with those acquired before the injection allowed for the manual outlining of the region of interest (ROI) encompassing the aortic lesions, thus facilitating the precise localization and volumetric assessment of the atherosclerotic plaques (pre-contrast shown in Figure 6A). This multimodal imaging approach provided a clear depiction of the scope and extent of atherosclerotic plaque within the vessels using SPECT and MR modalities.

**Figure 6.**
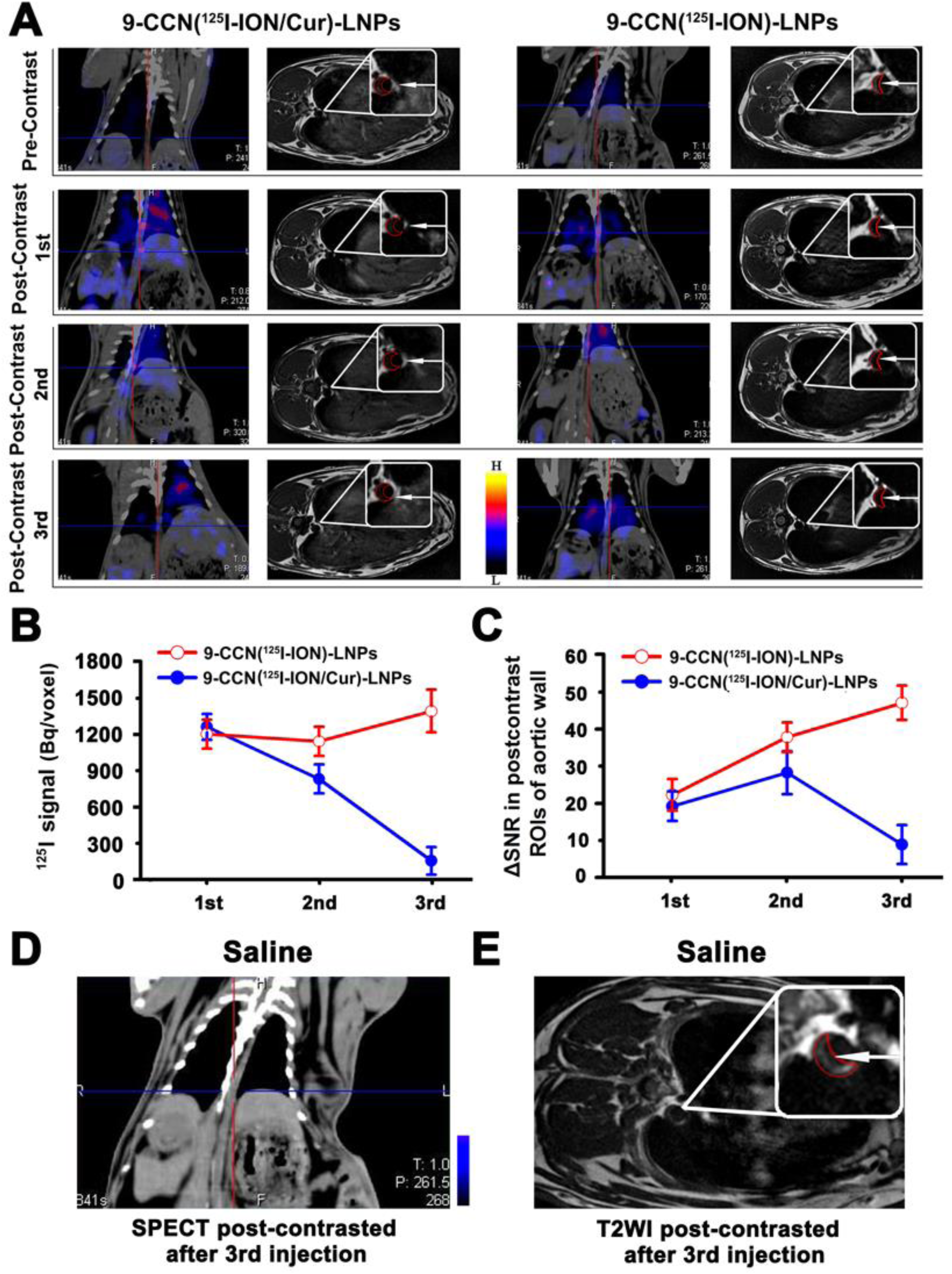
*In vivo* MRI and SPECT imaging for atherosclerosis plaques. New Zealand white rabbits (male, ∼3 kg) with AS were randomly administered with saline, 9-CCN[^125^I-ION/Cur]-LNPs or 9-CCN[^125^I-ION]-LNPs (three doses, 12.3 MBq/kg, ∼1.67 mg iron/kg, ∼17 mg curcumin/kg). (A) The Atherosclerotic plaque were monitored by SPECT and MRI scanning at 6 h and 36 h post-injection, respectively; (B) The radiotracer uptake (Bq/voxel) calculated by dividing radioactivity by volume from VOIs; (C) The SNR of the plaque using MRI measurement. (D) The plaque image on SPECT of the post-contrasted after 3^rd^ injection; (E) The plaque image on T2W of the post-contrasted after 3^rd^ injection. Data are expressed as mean ± SD (n = 3).

As a multimodal imaging probe for SPECT/MRI, careful consideration should be given to achieving a balanced combination of radionuclides and IONs to complement each other over both short and long terms. Although SPECT and MRI investigations were performed separately in this study, they contribute synergistically to the overall imaging performance. After the initial dose, no significant differences were observed between MLNPs and 9-CCN(^125^I-ION)-LNPS in terms of radioactivity and the change in SNR (ΔSNR) on MR images (Figure 6B&C). However, it was found that curcumin promoted the polarization of macrophages from an M1 to an M2 phenotype.

Overall, these findings underscore the potential of the multimodal imaging approach using 9-CCN[^125^I-ION]-NPs with curcumin for precise visualization and characterization of atherosclerotic plaque, while emphasizing the importance of a well-balanced and integrated SPECT/MRI strategy to enhance diagnostic capabilities. Thus, following the administration of the second dose, the radioactivity of MLNPs decreased from 1267 Bq/voxel observed after the first dose to 844 Bq/voxel, exhibiting a steady ^125^I signal that remained notably more prominent than that of 9-CCN(^125^I-ION)-LNPS. However, it is noteworthy that the △SNR of plaques in the MLNPs group showed no significant increase due to the rapid clearance and reduced uptake of IONs, as illustrated in Figure 6C. After the third dose, the ^125^I signals of MLNPs at the atherosclerotic plaque continued to diminish significantly to an almost non-detectable level, with a radioactivity measure of 165 Bq/voxel. In contrast, the average radioactivity of the ascending aorta treated with targeted lipid nanoparticles without the drug increased to 1409 Bq/voxel (Figure 6B). As a control group for biological studies, rabbits with AS were treated with saline. The SPECT and MR imaging of the pre-contrast condition mirrored the images obtained after the contrast agent injection. The plaque images on SPECT and T2WI following the third injection were depicted in Figure 6D&E, respectively.

The planar view of the radioactivity distribution provides a direct reflection of the atherosclerotic plaque composition, predominantly consisting of M1 macrophages, thus offering closer correlation with plaque vulnerability. Moreover, this planar imaging modality enables the enhancement of SNR, facilitating precise identification of small foci with low uptake and minimizing interference from the surrounding tissue exhibiting high radioactivity accumulation. In the planar images, radioactivity was observed to be distributed along the aorta in rabbits treated with 9-CCN(^125^I-ION/Cur)-LNPs, but nearly 90% lower compared to rabbits treated with 9-CCN(^125^I-ION)-LNPS. Additionally, in the vulnerable plaque treated with 9-CCN(^125^I-ION/Cur)-LNPs, the area of the red O-stained region was significantly reduced (Figure S3). This decrease could be attributed to the ability of curcumin to significantly lower lipid concentration by reducing atherogenic lipids, such as total cholesterol and LDL-C, in animal models and human subjects [20].

At 36 hours post-injection of the third dose, the △SNR of 9-CCN(^125^I-ION)-LNPS continued to increase, while that of MLNPs decreased to 47.5 due to ION absorption. The △SNR of MLNPs dropped to 9.1, lower than that observed after the first dose, suggesting that the uptake of ION in LNPs by M2 macrophages and the metabolism of retained ION in the plaque no longer led to a T2W signal, indicating a stabilization of the plaque. This observation was further corroborated by the iron distribution analysis of ex vivo plaques using Prussian blue staining after the third MR imaging, where the iron content in the plaques treated with MLNPs was remarkably lower than that in the LNPs without drugs (Figure S4).

In vivo MRI assessments revealed that MLNPs exhibited superior imaging capabilities, as evidenced by significantly increased accumulation of IONs in the 9-CCN(^125^I-ION/Cur)-LNP-treated group, as revealed by Prussian blue staining. Notably, apart from its enhanced diagnostic potential, MLNPs demonstrated superior therapeutic efficacy against AS plaques, as manifested by the substantial regression of atherosclerotic plaque in the 9-CCN(^125^I-ION/Cur)-LNP-treated group, with no observed plaque rupture. Conversely, in the 9-CCN(^125^I-ION)-LNP-treated groups, the plaque was still apparent.

As depicted in Figure 6, the radioactivity of MLNPs and 9-CCN(^125^I-ION)-LNPS exhibited significant differences at various stages of drug administration. After the first dose, the radioactivity of MLNPs and 9-CCN(^125^I-ION)-LNPS did not display notable differences due to the similar endocytosis activity of pro-inflammatory M1 macrophages. However, following the second and third doses, the radioactivity of MLNPs decreased substantially compared to that of 9-CCN(^125^I-ION)-LNPS. This disparity can be attributed to curcumin treatment, which effectively induced M1-M2 macrophage polarization, leading to a significant reduction in the endocytosis activity of macrophages.

Regarding ΔSNR, due to the consistent endocytosis activity of pro-inflammatory M1 macrophages, the ΔSNR of 9-CCN(^125^I-ION)-LNPS progressively increased, while that of MLNPs remained stable, consistent with the M1-M2 polarization induced by curcumin treatment. In summary, the simultaneous detection and therapy of atherosclerotic plaque were successfully achieved by MLNPs, through the co-delivery of a hybrid imaging agent and curcumin to macrophages. The therapeutic efficacy of MLNPs was accurately and dynamically visualized through radiography and MR imaging.

### 2.6. MLNPs induces M2 macrophage polarization in vivo

H&E and immunohistochemical staining of the ex vivo thoracoabdominal aorta provided revealing insights into the characteristics of the Atherosclerotic plaque in different treatment groups. In the 9-CCN(^125^I-ION)-LNPS-treated group, the plaques were observed at the verge of rupture and were extensively infiltrated with M1 macrophages, without significant distribution of M2 macrophages. In stark contrast, the 9-CCN(^125^I-ION/Cur)-LNP-treated group exhibited no evident plaque rupture, and the quantity of M2 macrophages surpassed that of M1 macrophages, as confirmed through immunohistochemistry and immunofluorescence staining of CD16/32 and CD206 (Figure 7A).Subsequent flow cytometry analysis further corroborated these findings, demonstrating a remarkable 4.4-fold increase in the number of M2 macrophages (CD206+/F4/80+) in the MLNPs group compared to the 9-CCN(^125^I-ION)-LNPS group, as depicted in Figure 7B & C. Additionally, MLNPs substantially upregulated the expression of M2-related Arg1 and Mgl1 by 6.9 and 14.3-fold, respectively, while concurrently downregulating the expression of INOs, relative to the 9-CCN(^125^I-ION)-LNPS group (Figure 7D).

**Figure 7.**
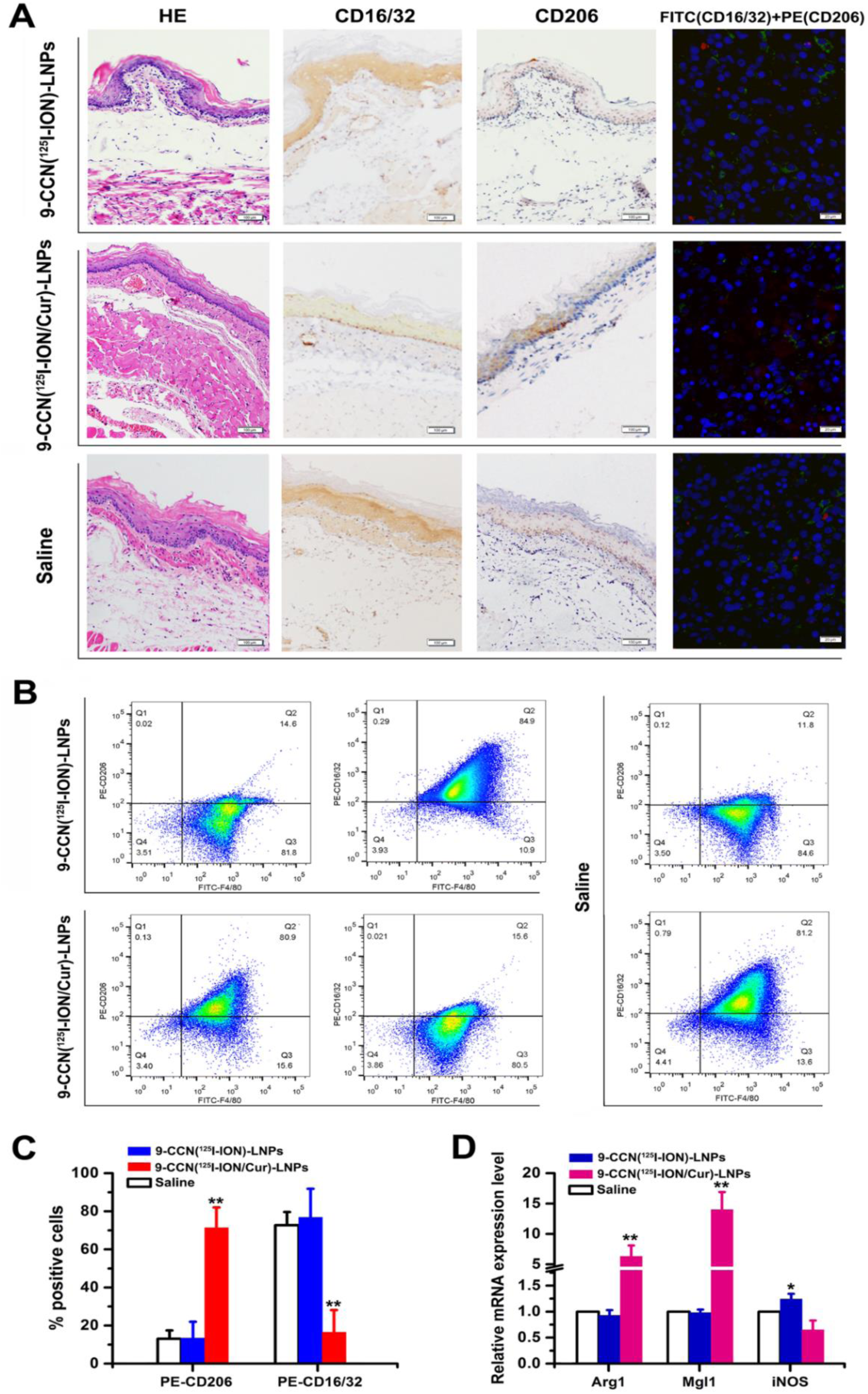
*Ex vivo* evaluation of atherosclerosis plaques of rabbit models. (A) Further confirmation of AS in the thoracoabdominal aortic of the rabbits by H&E staining. (B) The proportion of positively stained cells shown as scatter diagrams. A representative experiment was shown from three independent experiments. (C) The quantitative analysis of CD206+ and CD16/32+ macrophages in (B). (D) The mRNA levels of M1 and M2 macrophage markers (Arg1, Mgl1 and iNOS). The three or more groups are compared by one-way ANOVA with Dunnett’s post-test. *, p < 0.05; **, p < 0.01; ***, p < 0.001; n.s. represents not significant (p > = 0.05). Data are expressed as mean ±standard deviation (n = 3).

**Figure 8.**
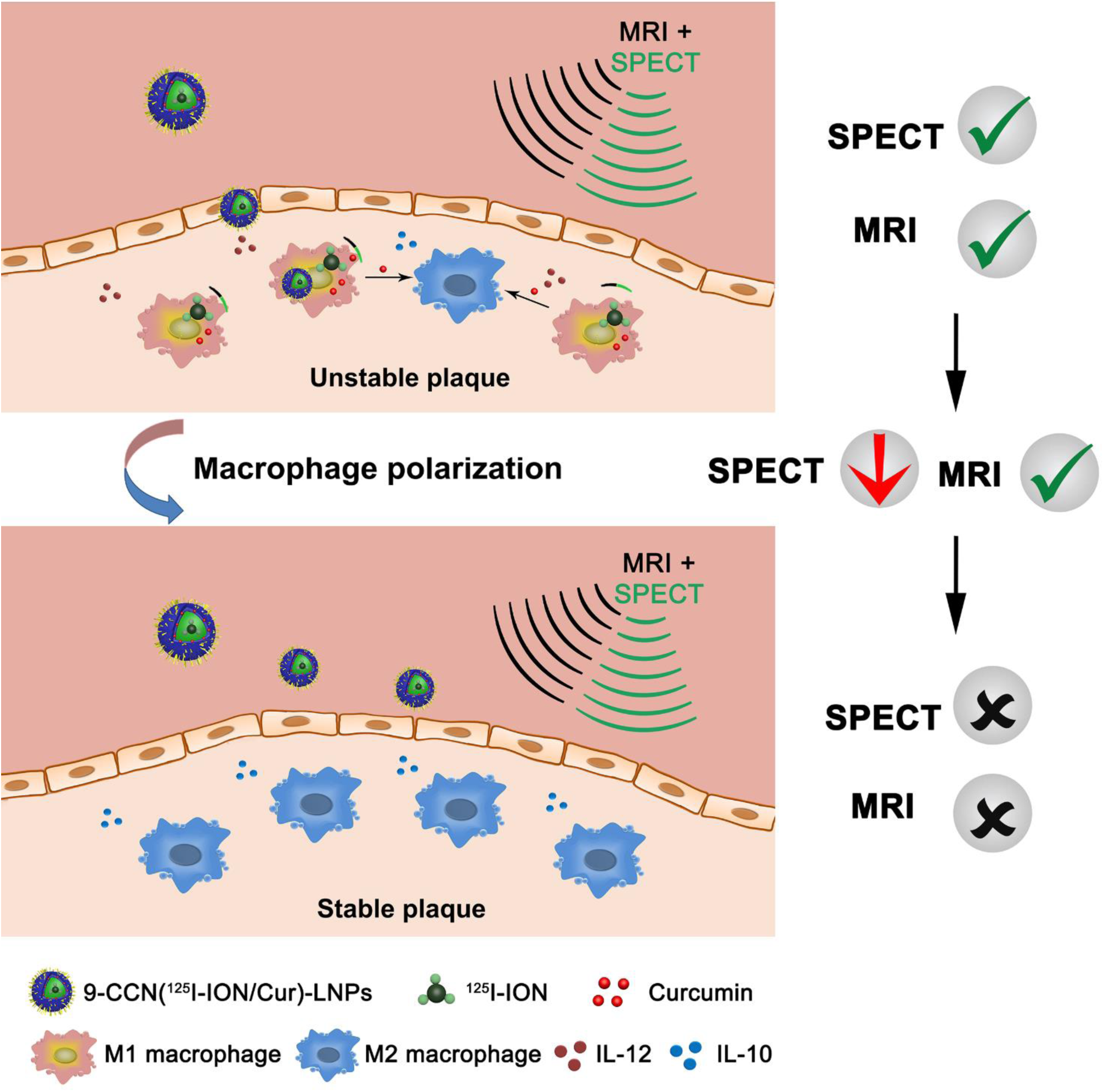
The theranostic mechanism of 9-CCN(^125^I-ION/Cur)-LNPs

These results collectively highlight the potent effect of 9-CCN(^125^I-ION/Cur)-LNPs in promoting M1-M2 macrophage polarization, thereby contributing to the stabilization of atherosclerotic plaque and mitigating the risk of rupture. Furthermore, the observed changes in M2-related gene expression further substantiate the immunohistochemical findings, establishing the therapeutic potential of MLNPs in promoting favorable macrophage phenotypes for the plaque management.

### 2.7. Safety and application prospects

The toxicity of the treatments was thoroughly assessed by conducting hematoxylin and eosin (H&E) staining on the excised organs from the rabbits (Figure S5). Notably, none of the rabbits exhibited any noticeable pathological changes in the H&E staining of all the examined organs. Moreover, the rabbits subjected to various treatments displayed no significant change in body weight when compared to the untreated control group (Figure S6). These findings collectively indicate that the rabbits demonstrated a commendable tolerance to all administered treatments, signifying their favorable safety profile.

## 3. Discussion

In this investigation, we have successfully engineered theranostic lipid-coated nanoparticles capable of co-delivering hybrid imaging agents, namely ^125^I-IONs and curcumin, for the dynamic imaging and treatment of atherosclerotic plaque. These multifunctional nanoparticles, which incorporated “eat-me” signals, were specifically designed to target inflammatory macrophages within vulnerable atherosclerotic plaque, subsequently releasing curcumin to promote the macrophage transformation from the M1 to M2 phenotype. Simultaneously, the progression of atherosclerotic plaque could be accurately probed using a multimodal imaging approach that combined the hypointensity observed on T2WI images with the predominantly radioactive accumulation seen on SPECT images.

The combined therapy utilizing macrophage-modulating curcumin and 9-CCN(^125^I-ION/Cur)-LNPs with the “eat-me” significantly enhanced the transformation rate from M1 to M2 phenotype, effectively reducing the risk of plaque rupture in vulnerable atherosclerotic plaque. The lipid-based nanoparticles were engineered to contain a hydrophobic core loaded with curcumin and a hydrophilic core loaded with the hybrid imaging agent ^125^I-IONs. The presence of the “eat-me” signal was crucial in maintaining the targeting activity of our 9-CCN(^125^I-ION/Cur)-LNPs towards macrophages.

The data presented above confirmed that 9-CCN(^125^I-ION/Cur)-LNPs efficiently delivered their payload to pro-inflammatory M1 macrophages through the “eat-me” signal. Once effectively targeted to M1 macrophages, these nanoparticles were readily internalized, releasing IONs, ^125^I, and curcumin into the cytoplasm. This process results in the downregulation of M1 markers and the upregulation of M2 markers. Both IONs and curcumin have been widely recognized for their respective roles in imaging and therapeutic applications. Curcumin, which is known for its cholesterol-lowering effects, has been reported to modulate macrophage polarization towards the M2 phenotype by inhibiting M1 markers and upregulating M2 markers.

In this study, we employed multimodal imaging combining SPECT and MRI techniques to accurately inspect and monitor the stabilization of macrophages from the M1 to M2 phenotype. While MRI offers high resolution, its sensitivity to smaller plaques may be limited. Integrating nuclear medical imaging, which uses radioactive materials for diagnosing various diseases, with MRI can address this sensitivity concern, combining the high specificity of functional imaging with the high resolution of structural imaging. We have previously used IONs for accurate and high-resolution localization of different liver tumors simultaneously. Additionally, our previous work reported the successful application of nanoparticles consisting of ^99^mTC and IONs for SPECT and MR imaging, allowing comprehensive assessment of atherosclerosis progression.

The combination of short-term radioactive reagents (using ^125^I in this study) and long-term ION-based contrast agents provided dynamic and comprehensive monitoring of the targeted delivery of curcumin to vulnerable atherosclerotic lesions and the theranostic procedure for atherosclerotic lesions. The use of the “eat-me” signal further facilitated targeted delivery to vulnerable plaques predominantly consisting of M1 macrophages, with limited distribution in stable plaques.

Regarding safety, 9-CCN(^125^I-ION/Cur)-LNPs did not exhibit severe toxicity to important organs in rabbits. This finding indicates that the use of these nanoparticles is expected to be safe in clinical applications, although additional safety studies are warranted. Furthermore, the local delivery of curcumin directly targeting plaques minimizes unpredictable absorption in other organs and tissues, thereby reducing potential side effects associated with systemic therapeutic drugs.

The theranostic mechanism of 9-CCN(^125^I-ION/Cur)-LNPs is attributed to their hydrophilic PEGylated lipid coating, which enables extended circulation time after intravenous injection. The presence of the “eat-me” signal in these nanoparticles allows specific binding to pro-inflammatory M1 macrophages within atherosclerotic plaques. Subsequent release of the hybrid imaging agent and curcumin results in facile radiography, MRI, and the desired polarization from M1 to M2 of macrophages. The M1-M2 polarization leads to the transformation of unstable atherosclerotic plaque into stable plaque, thereby reducing nanoparticle uptake, radioactivity, and achieving reliable MRI imaging.

As the results of this study have shown promising performance, further verification in human subjects, standardization of 9-CCN(^125^I-ION/Cur)-LNPs preparation, and rigorous assessment of metabolic and circulatory toxicity issues are essential steps in the forthcoming R&D period. These endeavors will pave the way for potential clinical applications of this innovative theranostic approach.

## 4. Conclusion

The current clinical landscape demands the development of theranostic agents that seamlessly integrate targeted therapy and contrast agents, enabling a dual-functional approach for atherosclerosis nuclear and MR imaging within a single dosage, while leading to the transformation of unstable atherosclerotic plaque into stable plaque. This study highlights the potential of 9-CCN(^125^I-ION/Cur)-LNPs, a novel formulation embodying unique targeting strategies, coalescing non-specific targeting with the therapeutic agent curcumin, to effectively investigate and address atherosclerotic plaque in vivo, utilizing both MRI and SPECT, despite their significant differences in sensitivity. To the best of our knowledge, this study represents the first demonstration of multifunctional lipid nanoparticles proficiently accomplishing simultaneous targeted therapy and highly contrast-enhanced SPECT/MRI tumor imaging via a single-dose administration, thus establishing a robust foundation for forthcoming clinical trials and immediate applicability within a clinical context.

## Author Contributions

Z. Shi, J. Huang and C. Chen contributed to the study concept and design, analysis, and interpretation of data, drafting/revising the manuscript for content, and statistical analysis. C. Chen participated in study concept and design as well as acquisition and analysis of nuclear medicine. X. Zhang was involved in analysis and interpretation of data. ZQ. Ma and Q. Liu contributed to the study concept and design, acquisition, analysis, and interpretation of data, and drafting/revising the manuscript for content. All authors have given approval to the final version of the manuscript.

## Acknowledgement

We gratefully thanks for the help of both Dr. Jie Gao and Mr. Huaiwen Chen due to drafting/revising the manuscript for content.

